# Epigenome-wide association study meta-analysis of wellbeing

**DOI:** 10.1101/2025.09.08.674802

**Authors:** Meike Bartels, Margot P. van de Weijer, Natalia Azcona-Granada, Bart M.L. Baselmans, Matthew Suderman, Mette Soerensen, Jadwiga Buchwald, Rosa H. Mulder, Rajesh Rawal, Michelle Luciano, Marc Jan Bonder, Priyanka Choudhary, Estelle Lowry, Penelope A. Lind, Joel Schwartz, Birgit Debrabant, Miina Ollikainen, Janine F. Felix, Marian J. Bakermans-Kranenburg, Henning Tiemeier, Christian Gieger, Melanie Waldenberger, Nicholas G. Martin, Pantel Vokonas, Andrea A. Baccarelli, Kaare Christensen, Jaakko Kaprio, Marinus H. van IJzendoorn, Rebecca T. Emeny, Ian J. Deary, Lude Franke, Sylvain Sebert, Allan F. McRae, Avron Spiro, Jenny van Dongen

**Author notes:** Corresponding author: Dr. Jenny van Dongen Department of Biological Psychology Vrije Universiteit Amsterdam Van Der Boechorststraat 7 1081 BT Amsterdam The Netherlands.

## Abstract

Wellbeing is associated with both behavioral phenotypes as well as several key life outcomes, such as health, employment, and coping with stressful events. These phenotypes associated with wellbeing could be potential indicators of differential epigenetic patterns between individuals that differ in their levels of wellbeing. We performed the largest epigenome-wide (EWAS) meta-analysis of wellbeing to date by combining whole blood DNA methylation data (Illumina 450k array) from 13 cohorts from Europe, Australia, and the USA (*N* = 10,757 participants). After correcting for smoking and BMI, no epigenome-wide significant methylation sites were identified. We tested whether a weighted methylation score (MS) based on leave-one-cohort-out EWAS meta-analysis summary statistics predicted wellbeing in an independent cohort, and whether prediction was significant over and above the polygenic score (PGS) for wellbeing. The MS was associated with wellbeing (variance explained=0.22%, *p*=0.03) and was no longer significant after adding the polygenic score (PGS; variance explained=0.43%, *p*=0.0046, MS; variance explained=0.07%, *p*=0.2842). We further compared DNA methylation levels in 16 pairs of monozygotic twins discordant for wellbeing. These analyses revealed no significant within-pair DNA methylation differences at the top-sites from the meta-analysis or in MS. Our results suggest that larger EWAS meta-analyses with uniform phenotype assessment are required to identify methylation sites associated with wellbeing.

## Introduction

There has been progress in the identification of genetic variants influencing individual differences in wellbeing^1–4^. For example, the most recent meta-analysis by Baselmans and colleagues identified over 300 loci jointly explaining about 1% of the phenotypic variation^4^. The next challenge is to improve our understanding of the biological effects of these genetic susceptibility loci, especially since the actual causal genes are not necessarily proximal to the lead SNPs identified in genome-wide association studies (GWASs). Supported by the observation that GWAS variants are preferentially located in enhancers and open chromatin regions^5,6^, the majority of common genetic risk factors are predicted to influence gene regulation through effects on epigenetic marks such as DNA methylation rather than directly affecting the coding sequence of transcribed proteins^7^. We previously applied an MWAS (methylome-wide association study) approach which combined information from methylation Quantitative Trait Loci (mQTLs) and GWAS summary statistics of wellbeing to identify SNPs whose effect on wellbeing was mediated by DNA methylation^4^. This approach identified 913 CpGs whose methylation level mediated the effect of *cis* genetic variants on wellbeing^4^.

Accumulating lines of evidence point towards a key role for epigenetic mechanisms regulating gene expression during brain development, maturation, and learning^8^. Besides playing an important role in the development of the brain, DNA methylation has been linked to various health outcomes that are associated with wellbeing, such as obesity^9^, type 2 diabetes^10,11^, and depression^12,13^. Variation in DNA methylation between individuals can be the result of environmental influences^14–16^. For instance, various early life exposures such as childhood abuse^17^, and prenatal exposure to maternal famine^18^ can induce long-term changes in DNA methylation^17–19^. Additionally, methylation variation can result from exposures later in life. Cigarette smoking is perhaps the best studied example in human epigenome-wide association studies (EWASs) and has been associated with differential methylation at thousands of CpG sites^20,21^.

Wellbeing is linked to numerous exposures and behaviors across the life course, such as income and employment, health, neighborhood environment (e.g., neighborhood safety), smoking, stress, alcohol use, and several social factors such as friendship patterns^22^. These exposures and behavioral characteristics are potential candidate drivers of differential epigenetic patterns between individuals reporting higher or lower levels of wellbeing. We previously investigated the association between DNA methylation in blood and wellbeing in the Netherlands Twin Register (NTR, *N* = 2,519) using an EWAS approach, reporting two CpG sites that showed a statistically significant association after Bonferroni correction^23^. Gene ontology (GO) analysis highlighted enrichment of several central nervous system categories among methylation sites with the smallest p-values. However, replication of these results in larger samples is warranted.

Here, we performed a large (*N* = 10,757) EWAS meta-analysis of whole blood DNA methylation data (Illumina 450K array) from 13 cohorts with various measures of wellbeing. In the NTR cohort, we subsequently assessed the predictive value of wellbeing DNA methylation scores (MS) created using weights from a leave-one-out meta-analysis without NTR. We also tested if the MS predicted wellbeing over and above the polygenic score (PGS) for wellbeing, to examine if PGS and MS capture independent information about wellbeing. Finally, in a unique group of monozygotic (MZ) twins from pairs discordant for wellbeing, we exploratively tested for differentially methylated CpG sites, ruling out possible confounding genetic -and environmental factors.

## Methods

### Epigenome-wide association study

Data on wellbeing, white blood cell counts, and DNA methylation levels were available for 13 cohorts (*N* = 10,757) of which 10 cohorts (*N* = 8,675) also had information on the covariates body mass index (BMI) and smoking status (see **Table 1**). The included cohorts were from Europe, Australia, and the United States (European ancestry). All participants provided written informed consent regarding participation in the individual studies, and all studies were approved by the local research ethics committees or institutional review boards. More information on the individual cohorts can be found in **Supplementary Tables 1**.

**Table 1:**
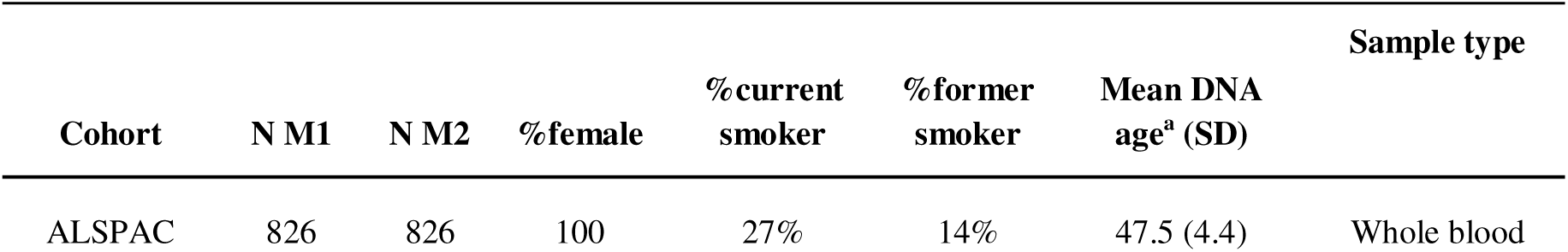
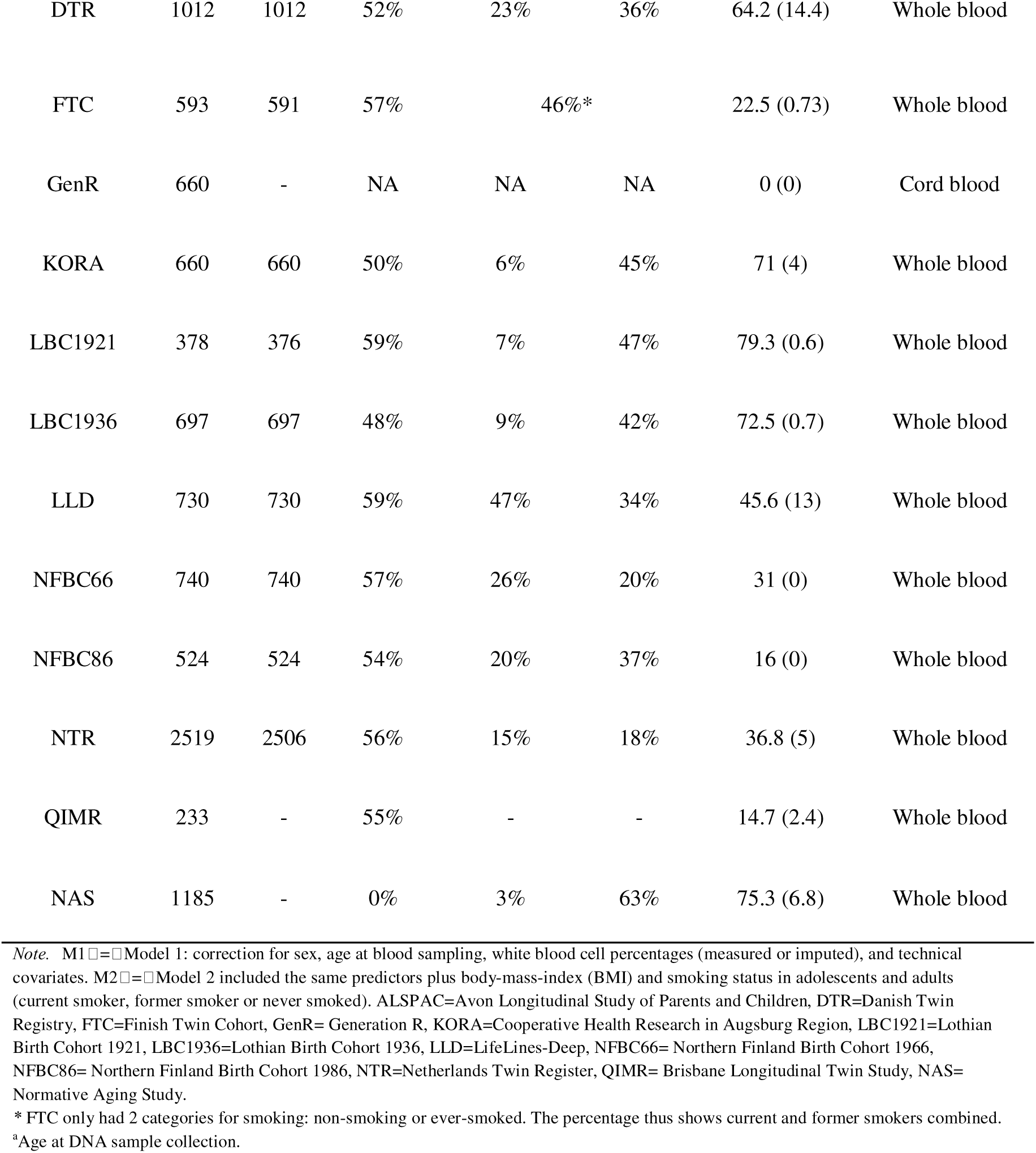
Cohort characteristics

| Cohort | N M1 | N M2 | %female | %current smoker | %former smoker | Mean DNA age<sup>a</sup> (SD) | Sample type |
| --- | --- | --- | --- | --- | --- | --- | --- |
| ALSPAC | 826 | 826 | 100 | 27% | 14% | 47.5 (4.4) | Whole blood |
| DTR | 1012 | 1012 | 52% | 23% | 36% | 64.2 (14.4) | Whole blood |
| FTC | 593 | 591 | 57% | 46%* |  | 22.5 (0.73) | Whole blood |
| GenR | 660 | - | NA | NA | NA | 0 (0) | Cord blood |
| KORA | 660 | 660 | 50% | 6% | 45% | 71 (4) | Whole blood |
| LBC1921 | 378 | 376 | 59% | 7% | 47% | 79.3 (0.6) | Whole blood |
| LBC1936 | 697 | 697 | 48% | 9% | 42% | 72.5 (0.7) | Whole blood |
| LLD | 730 | 730 | 59% | 47% | 34% | 45.6 (13) | Whole blood |
| NFBC66 | 740 | 740 | 57% | 26% | 20% | 31 (0) | Whole blood |
| NFBC86 | 524 | 524 | 54% | 20% | 37% | 16 (0) | Whole blood |
| NTR | 2519 | 2506 | 56% | 15% | 18% | 36.8 (5) | Whole blood |
| QIMR | 233 | - | 55% | - | - | 14.7 (2.4) | Whole blood |
| NAS | 1185 | - | 0% | 3% | 63% | 75.3 (6.8) | Whole blood |

*Note.* M1 = Model 1: correction for sex, age at blood sampling, white blood cell percentages (measured or imputed), and technical covariates. M2 = Model 2 included the same predictors plus body-mass-index (BMI) and smoking status in adolescents and adults (current smoker, former smoker or never smoked). ALSPAC=Avon Longitudinal Study of Parents and Children, DTR=Danish Twin Registry, FTC=Finish Twin Cohort, GenR= Generation R, KORA=Cooperative Health Research in Augsburg Region, LBC1921=Lothian Birth Cohort 1921, LBC1936=Lothian Birth Cohort 1936, LLD=LifeLines-Deep, NFBC66= Northern Finland Birth Cohort 1966, NFBC86= Northern Finland Birth Cohort 1986, NTR=Netherlands Twin Register, QIMR= Brisbane Longitudinal Twin Study, NAS= Normative Aging Study.
\* FTC only had 2 categories for smoking: non-smoking or ever-smoked. The percentage thus shows current and former smokers combined.
<sup>a</sup>Age at DNA sample collection.

### Wellbeing Measurements

Detailed information on the wellbeing measures for each cohort are provided in **Supplementary Table 2**. All items (single or summed) were wellbeing questions, except for Lifelines (LLD). LLD derived a sumscore from the positive-affect negative-affect (PANAS) questionnaire ^24^ with items such as ‘interested’, ‘enthusiastic’, ‘proud’ and ‘inspired’.

### DNA Methylation BeadChips

DNA methylation was assessed with the Illumina HumanMethylation450 BeadChip (450k array) in whole or cord blood. All cohorts analyzed DNA methylation β-values, which range from 0 to 1, indicating the proportion of DNA that is methylated at a CpG in a sample. More information on the methylation data of the individual cohorts is presented in **Supplementary Table 3**.

### Epigenome-wide Association Analysis

EWAS analyses were performed according to a standard operating procedure (See **Supplementary Information 1**). In each cohort, the association between DNA methylation level and wellbeing was tested under a linear model with DNA methylation as outcome, and correction for relatedness of individuals, using the r package generalized estimation equations (gee), where applicable (see **Supplementary Table 4**). Gee corrects for familial relatedness by using an exchangeable conditional covariance matrix and basing the tests on robust (sandwich corrected) standard errors. Two models were fitted. The first model included the following predictors: wellbeing, sex (if applicable), age at blood sampling (if applicable), the square of the Z-score of age at blood sampling (hereafter referred to as standardized(age at blood sampling)^2^), white blood cell percentages (measured or imputed), and cohort-specific technical covariates (such as sample plate and array row). The second model additionally included BMI and smoking status (current smoker/former smoker /never smoked or ever smoker/never smoker). We adjusted for BMI and smoking because they are known to have large effects on DNA methylation and can be confounders^20,25^

### Quality Control

The following methylation sites were removed from the meta-analysis: methylation sites on the sex chromosomes, methylation sites with more than 5% missing data in a cohort, methylation sites interrogated by probes overlapping with SNPs with a minor allele frequency (MAF) > 0.01 in the 1000G EU population or GONL population that affect the CpG or single base extension site ^26^, and ambiguously mapping probes reported by Chen *et al* with an overlap of at least 47 bases per probe^27^. In addition, we applied a minimum sample size filter (N > 10000 for model 1), to remove probes that were missing in multiple cohorts. The number of probes after filtering was 387188. Detailed cohort-level quality control and filtering of the methylation data is described in **Supplementary Table 3**. The R package Bacon was used to compute the Bayesian inflation factor and to obtain bias-and inflation-corrected test statistics prior to meta-analysis ^28^.

### Meta-analysis

Fixed-effects meta-analyses were performed in METAL^29^. We used the p-value-based (sample size-weighted) method because the measurement scale of wellbeing measures differs across studies. The I^2^ statistic from METAL was used to describe heterogeneity. Statistical significance was assessed considering Bonferroni correction for the number of sites tested (α=1.3×10^-7^). Additionally, we considered the False Discovery Rate (5%). False discovery rate (FDR) q-values were computed with the R package q-value with default settings.

### Methylation Score

Based on summary statistics from the whole blood EWAS meta-analysis of wellbeing, DNA methylation scores (MS) were created in the adult NTR cohort (whole blood 450k array data) to test the combined predictive value of DNA methylation sites associated with wellbeing in whole blood^30,31^. For this purpose, the meta-analysis of EWAS model 2 was rerun after excluding NTR (new *N*=6156). We applied a minimum sample size filter (N > 6000 for model 2) to remove probes with high missingness. For each NTR participant, a weighted sum score was calculated by multiplying the methylation value at a given CpG by the effect size (z-score) from the model 2 EWAS meta-analysis, and then summing these values over all genome-wide CpGs:

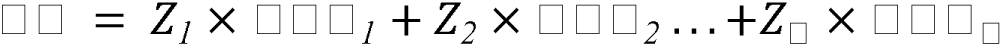

Where MS=Methylation score, CpG_i_ is the methylation level at CpG site i in the NTR participant, and Z_i_ is the effect size (z-score) at CpGi from the EWAS meta-analysis without NTR. We compared the performance of DNA methylation scores generated with pruning (removing CpGs with a correlation > 0.1) or without pruning, as previously described in Odintsova et al 2021^31^, and compared scores generated with various subsets of CpGs based on meta-analysis different p-value thresholds: p < 1 (all CpGs), p < 1 × 10^−1^, p < 1 × 10^−2^, p < 1 × 10^−3^ , p < 1 × 10^−4^ and p < 1 × 10^−5^. Pruning is performed to remove correlated CpGs that are redundant (and potentially add noise to scores). The expectation is that if the set of CpGs associated with a trait is correlated (and especially if correlations are strong or abundant), pruning will improve performance of the MS.

Wellbeing scores were regressed on the MSs with gee models, which were fitted with the R package ‘gee’. The following settings were used: Gaussian link function (for continuous data), 100 iterations, and the ‘exchangeable’ option to account for the correlation structure within families and within persons. Wellbeing was regressed on each MS plus the same covariates as included in EWAS model 2; white blood cell counts, sample plate (dummy-coded), array row, age at blood sampling, standardized (age at blood sampling)^2^, sex, BMI, and smoking (current smoker/former smoker /never smoked) as covariates. In each analysis, MSs and wellbeing scores were standardized, that is, the mean was subtracted from each score and the remaining value was then divided by the standard deviation. This makes the estimated regression coefficient for the effect of the predictor MS on the outcome wellbeing equivalent to a (semipartial) correlation. The proportion of variance in wellbeing explained by the MSs, after accounting for covariates, can subsequently be obtained by squaring the regression coefficient. This value was multiplied by 100 to obtain the percentage of variance explained.

### Combined analysis of a polygenic score and methylation score

Next, we took the best-performing MS and tested whether the MS significantly predicted wellbeing over and above a polygenic score (PGS) based on GWAS summary statistics. For this purpose, we used the N-weighted genome-wide association meta-analysis (NGWAMA) from Baselmans et al.^4^ excluding samples from the Netherlands Twin Register. Using these summary statistics, polygenic scores were created using the following strategy.

The weights used for the polygenic scores were based on the wellbeing spectrum multivariate NGWAMA including the highly genetically correlated measures of life satisfaction, positive affect, depression, and neuroticism. Scores were based on the intersection of SNPs available in any of these GWAMAs. Quality control parameters for genotype data in NTR are described in **Supplementary Table 5**. In total, 1,224,793 SNPs passed QC and were used to construct polygenic scores. Before aggregating the individual effect sizes into polygenic scores using plink^32^, we adjusted the effect sizes for linkage disequilibrium (LD) using LDpred^33^. LPpred requires to specify the fraction of SNPs with a non-zero effect on wellbeing, which was set to 1. The prediction of wellbeing by PGS and MS was tested with the R package ‘gee’. We corrected for the same covariates as in the EWAS (that is: age at blood sampling, standardized (age at blood sampling)^2^, sex, BMI, smoking (current smoker/former smoker /never smoked), sample plate (dummy-coded), array row, and percentages of monocytes, eosinophils, and neutrophils), plus the first 10 genomic principal components (PCs), and genotyping platform dummies.

### Discordant twin pair analysis

To control for genetic and shared environmental confounding we used the unique characteristics of monozygotic (MZ) twin pairs to help understand the association between DNA methylation and wellbeing. MZ co-twins share nearly 100% of their DNA sequence as well as their common environment during childhood and adolescence. Therefore, MZ twins from pairs discordant for wellbeing form the ideal case-control groups because they are perfectly matched on many possible confounding factors, including age, sex, genetic make-up, and numerous shared environmental factors^34^.

As the criterion for discordance, we selected twins who had a within-pair difference wellbeing score of >3 standard deviations (SDs) on the satisfaction with life scale. Based on this threshold, 16 MZ twin pairs out of 624 MZ twin pairs (2 %) from the NTR with data on wellbeing and DNA methylation were classified as discordant for wellbeing. The mean wellbeing score for the high-scoring twin (Mean_wellbeing_ = 30.62) was on average 19 points higher than that of the low-scoring twin (Mean_wellbeing_= 11.06) (see **Figure 1**), while the mean (sd) of the full sample was 27.07 (5.46) (See **Table 2**). Additionally, we identified twins from pairs concordant in their wellbeing score, i.e., with identical wellbeing scores (N= 91). From these concordant twin pairs, we randomly selected 16 concordant twin pairs (Mean_wellbeing_ = 29) to correspond to the number of discordant twin pairs. The data from concordant pairs were used to compare the distribution of within-pair DNA methylation differences.

**Figure 1:**
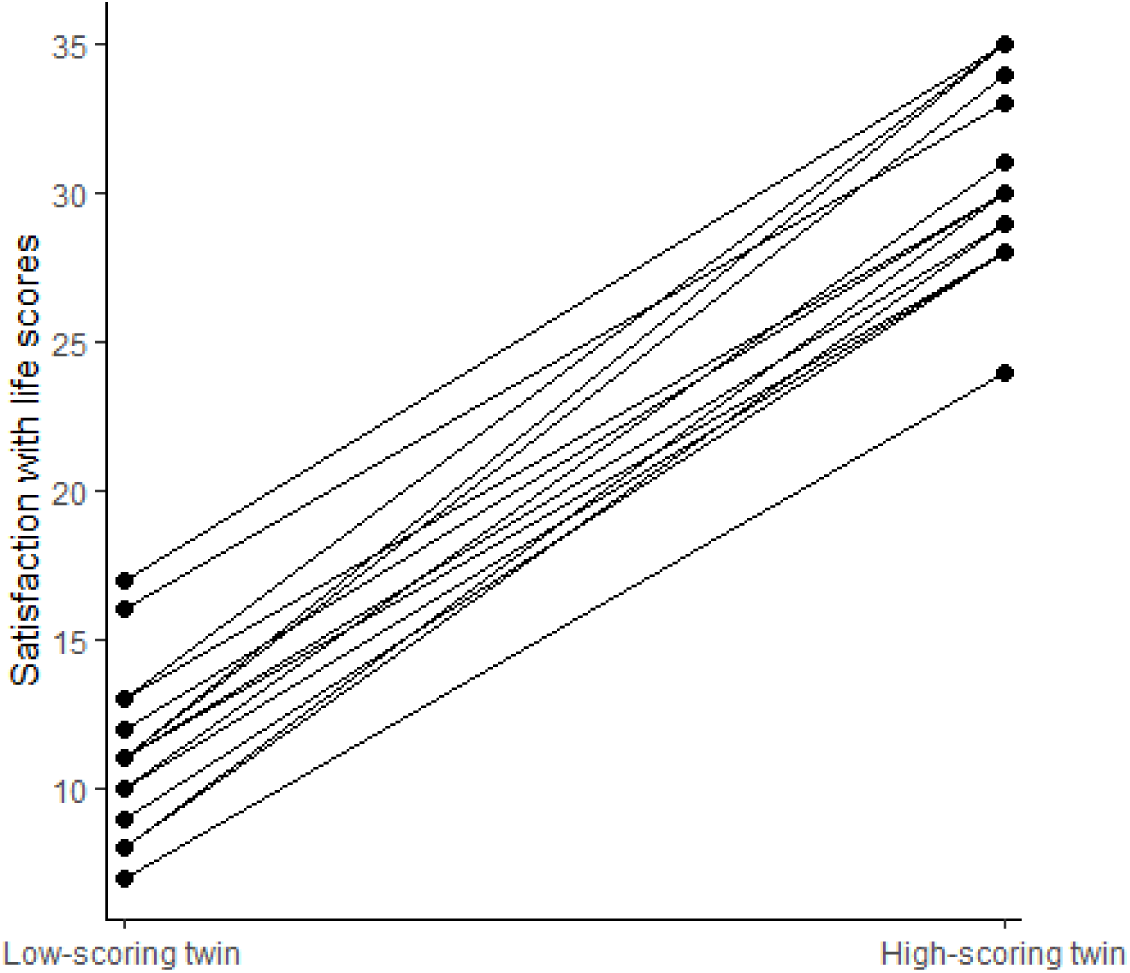
Satisfaction with life scores for monozygotic twin pairs discordant for wellbeing (N=16)

**Table 2:** Descriptives wellbeing (WB) discordant MZ twins

|  | Low WB twin | High WB twin |
| --- | --- | --- |
| Mean WB (SD) | 11.06 (2.72) | 30.63 (3.11) |
| WB range | 7-17 | 24-35 |
| *N* males/females | 1/15 | 1/15 |
| Mean age (SD)<sup>a</sup> | 36.51(11.59) | 36.36 (11.69) |
| Mean BMI (SD) | 24.31 (5.15) | 24.96 (5.15) |
| Neutrophil (%) | 51.18 (9.34) | 53.34 (10.26) |
| Monocyte (%) | 7.86 (1.81) | 8.05 (2.24) |
| Eosinophil (%) | 2.74 (1.67) | 3.03 (2.24) |
| *N* non-smoker/former<br>smoker/current smoker | 13/3/0 | 11/4/1 |

WB=wellbeing. <sup>a</sup>Age at DNA sample collection. Note: in the Netherlands Twin Register, participants were visited at home for collection of blood samples. Minor differences in age at sample collection reflect that co-twins could not always be visited on the same day.

We compared the similarity in methylation levels in discordant MZ pairs and concordant MZ twin pairs. DNA methylation levels were residualized for BMI, smoking, sample plate (dummy coded), array row, and percentages of monocytes, eosinophils, and neutrophils in advance with a linear regression model (i.e., the same covariates as included in the EWAS minus age and sex because MZ twins have the same age and sex and the blood sample was collected at the same age). The residuals were used as input for paired t-tests. We tested whether the top 10 CpG sites with the lowest p-value from the EWAS meta-analysis excluding NTR showed differences in DNA methylation in the discordant MZ pairs at a Bonferroni-corrected significance threshold of α= 0.05/10= .005, and tested if the overall distribution of the average absolute within-pair differences of residualized methylation beta-values differed between discordant and concordant twin pairs for the top 10 CpGs using a Kolmogorov-Smirnov test. Second, we tested if the DNA methylation score (for the pruned score with P<1) for wellbeing differed in discordant MZ pairs with a paired t-test.

## Results

### Peripheral blood meta-analyses

We performed two meta-analyses. The first EWAS meta-analysis (Model 1) included 13 studies with peripheral or cord blood DNA methylation data (N= 10,757) but was not corrected for smoking and BMI. The second EWAS meta-analysis (Model 2) consisted of 10 studies (N = 8,675) with peripheral blood DNA methylation data, as well as information about smoking data and BMI. The full summary statistics can be found in **Supplementary Tables 6-7**. Results for the top 10 sites from Model 2 are shown in **Table 3**. No methylation sites were significantly associated with wellbeing in model 1 (supplementary figure 1 and supplementary figure 2) or model 2 (figure 2 and 3) after Bonferroni correction or at a false discovery rate of 5%. When inspecting heterogeneity statistics for both models, we observed that the top 10 sites (based on the outcomes from Model 2) showed mixed degrees of between-study heterogeneity (Model 1: mean I^2^ = 12.89%, range=0-79.4%, Model 2: mean I^2^ = 12.99%, range = 0-81.9%), but only 2 CpGs (Model 1) and 1 CpG (Model 2), respectively, showed significant heterogeneity after Bonferroni correction (α=1.2×10^-7^). We repeated the meta-analyses of model 1 after exclusion of one cohort with cord blood DNA methylation data (New N= 10097). The correlation between effect sizes (Zscores) from these two meta-analyses was high (*r*=0.97). The meta-analysis without cord blood samples yielded no significant differentially methylated sites and showed a similar degree of heterogeneity (mean I^2^ =13.19%, range=0-81, significant heterogeneity at 1 CpG).

**Figure 2:**
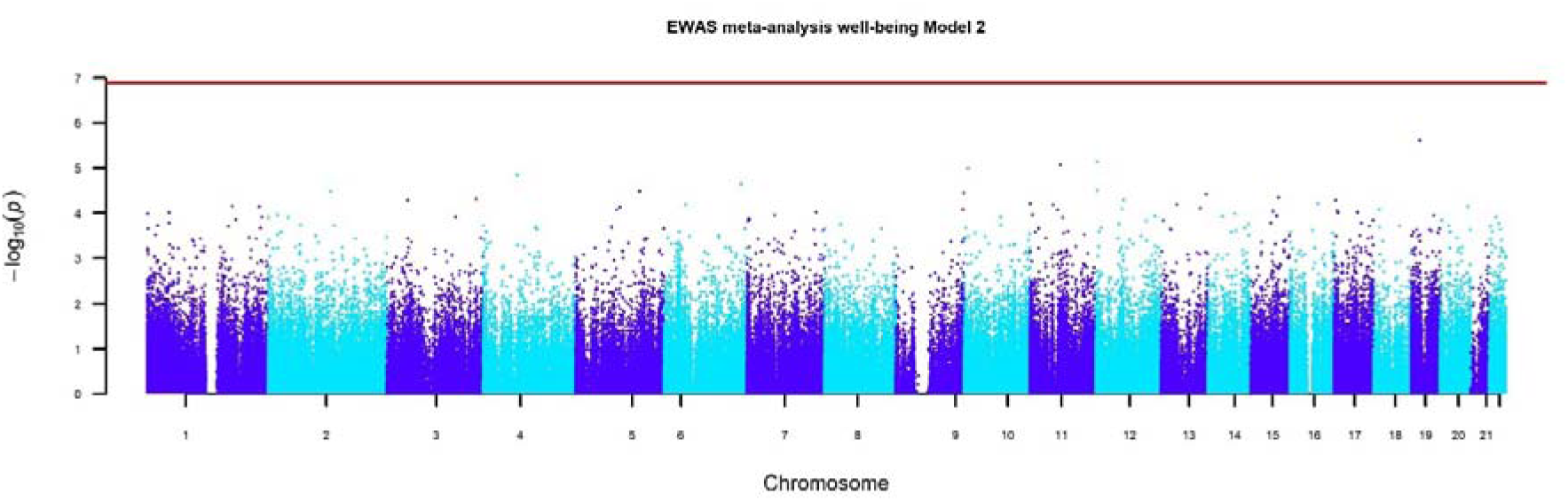
Manhattan plot for the EWAS meta-analysis with correction for smoking and BMI (model 2) The Manhattan plot shows the p-values for autosomal methylation sites from the EWAS meta-analysis of model 2, which included correction for smoking and BMI.

**Figure 3:**
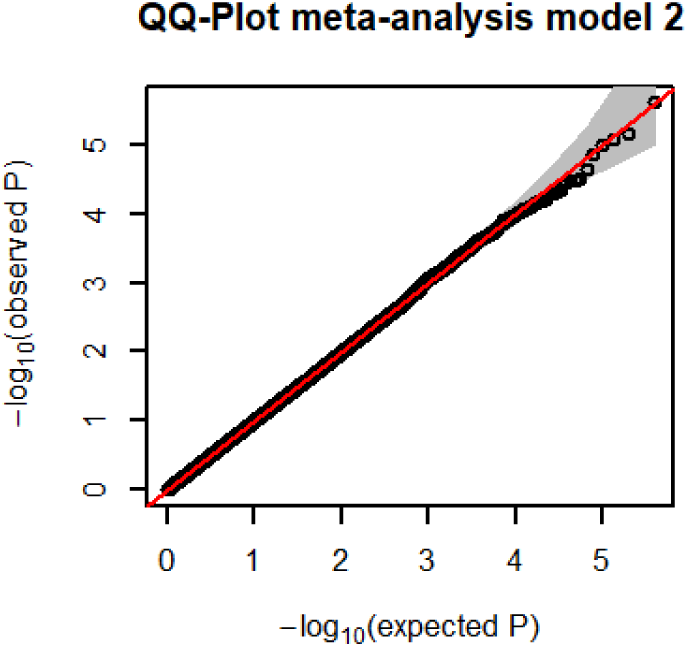
EWAS meta-analysis (model 2) QQ-Plot QQ-plot from the EWAS meta-analysis of model 2, which included correction for smoking and BMI. The inflation factor (lambda) was 0.99.

**Table 3:**
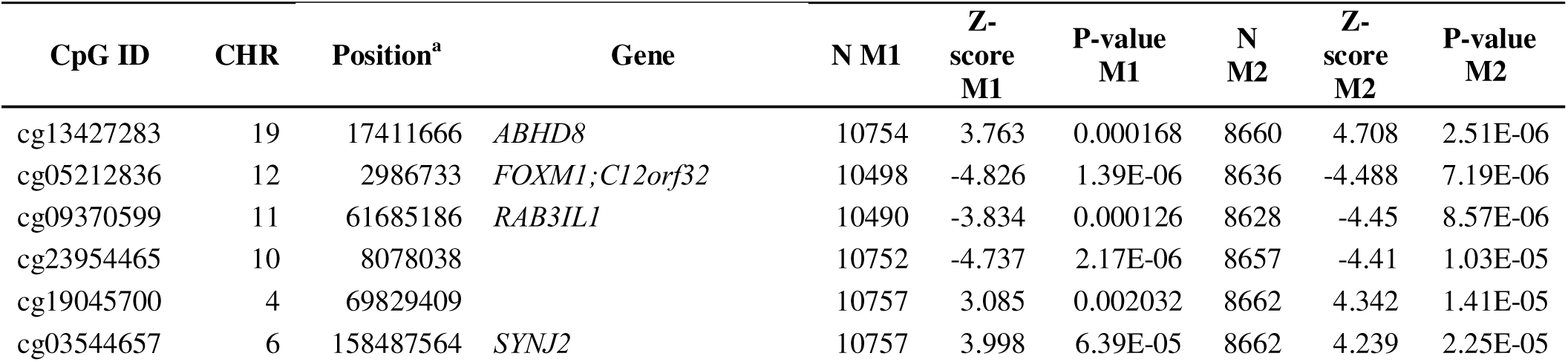
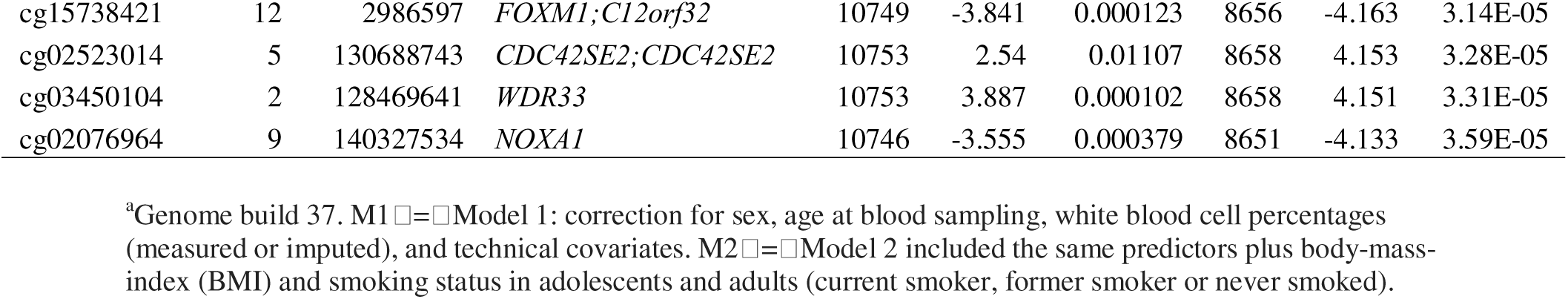
Top-sites associated with wellbeing from the EWAS meta-analysis of model 2

### DNA methylation scores

We computed weighted DNA methylation scores in the NTR (peripheral blood, *Mean*_age_ = 37.77 years, *SD* = 12.99, *N* = 2506), based on the leave-one-out EWAS meta-analysis of model 2 without NTR. The best-performing MS that positively predicted wellbeing (which was the pruned score, with a p-value threshold of P<1) explained 0.22% of the variation in wellbeing (b=0.047, p=0.0318), after adjusting for technical covariates, sex, age, standardized (age at blood sampling)^2^, cell counts, BMI, and smoking (**Supplementary Table 8**). Repeating the same analysis in a smaller subsample of NTR individuals who never smoked, the methylation score was no longer significantly predictive of wellbeing, but the effect size was fairly similar (*N* = 1437, *b*= 0.0357, *p* = 0.2357) suggesting that smoking does not drive the association between the methylation score and wellbeing.

### Combined performance of methylation and polygenic scores

Next, we tested whether the methylation score predicted wellbeing over and above a polygenic score (*N* = 2078). In a model that included the wellbeing polygenic score and the wellbeing methylation score, the wellbeing polygenic score was significant (*ß* = 0.0659, *R*^2^ = 0.4345%, *p* = 0.0046), but the methylation score was not significant (*ß* = 0.0256, *R*^2^ = 0.0653%, *p* = 0.2842, see **Supplementary Table 9**).

### Discordant twin pair analysis

We tested for differences in DNA methylation levels in monozygotic twin pairs discordant for wellbeing. There were no significant DNA methylation differences within the discordant pairs for the top 10 CpG sites from the EWAS meta-analysis without NTR (**Supplementary table 10**). Five of the 10 top CpGs from the meta-analysis without NTR showed the same direction of effect in discordant pairs. The Q-Q plot for the comparison of discordant monozygotic twin pairs based on all genome-wide CpG sites is shown in **Supplementary** Figure 3, illustrating that there was no inflation of test statistics and no genome-wide significant differences.

The distribution of the average within-pair absolute residual methylation difference for the top 10 CpGs did not differ significantly between discordant and concordant pairs (*p*=.99, Kolmogorov-Smirnov test) (see Figures 4 and 5). Additionally, the DNA methylation scores of the discordant twins did not differ significantly between the wellbeing-high and wellbeing-low scoring twins from discordant monozygotic twin pairs (t=.12, p=.91).

**Figure 4.**
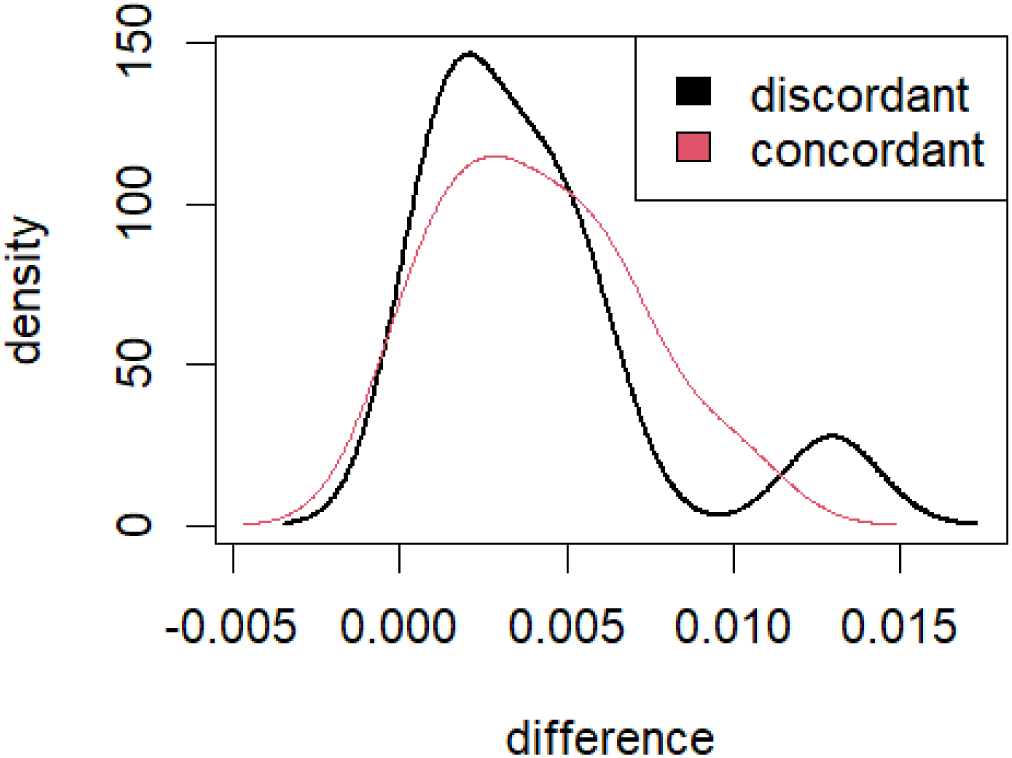
Density plot of the overall distribution of absolute within-pair DNA methylation differences in identical twin pairs based on the top 10 CpG sites from the EWAS meta-analysis of wellbeing

**Figure 5.**
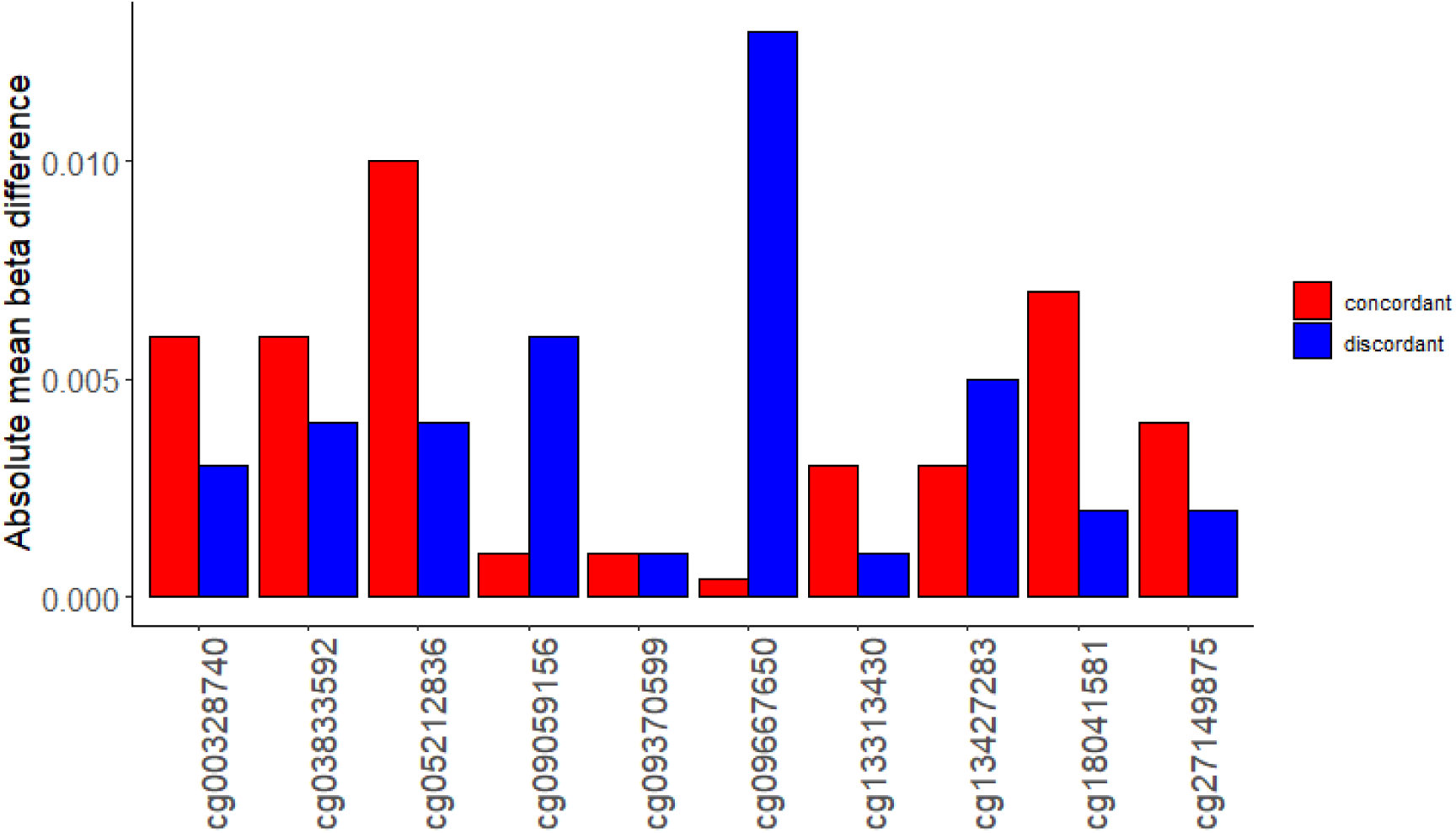
Absolute within-pair DNA methylation differences in wellbeing discordant and wellbeing concordant twin pairs for the top 10 CpGs from the EWAS meta-analysis excluding NTR. For each of the top 10 CpG sites from the leave-one-out meta-analysis without NTR (model 2), the absolute average within-pair difference in DNA methylation is plotted.

## Discussion

This study is the first epigenome-wide association study meta-analysis of wellbeing. No genome-wide significant methylation sites were identified either without (N = 10,757) or with correction for BMI and smoking (N = 8,675). A weighted DNA methylation score based on all genome-wide CpGs from the summary statistics from our meta-analysis of wellbeing explained ∼0.2% of the variance of wellbeing but was no longer significantly predictive of wellbeing after adding the PGS based on the largest GWAS of the wellbeing spectrum to the model^4^. In a secondary exploratory analysis, we used the unique characteristics of MZ twins to investigate site-specific methylation differences in monozygotic twin pairs discordant for wellbeing. This analysis did not identify significant methylation differences in wellbeing discordant monozygotic twin pairs.

Although our EWAS meta-analysis is among the largest EWAS meta-analyses today with a sample size of ∼11,000, it could be that the sample size is still too small to detect genome-wide significant CpGs for a complex behavioural trait such as wellbeing. A recent EWAS meta-analysis of aggressive behavior^35^, with a sample of over 14,000 individuals, also did not identify significant methylation sites after correcting for smoking and BMI. We found that DNA methylation scores for wellbeing explained less variation (∼0.2%) compared to methylation scores for, for instance, smoking (∼60%), BMI (∼10%), and educational attainment (∼2%)^31,36^; traits for which EWASs have been successful in identifying genome-wide significant CpGs with smaller sample sizes. Although the explained variance of the wellbeing methylation score is small, it is in the same range as the variance explained by wellbeing polygenic scores based on DNA markers^2,4^.

We performed follow-up analyses in a unique group of MZ twin pairs discordant for wellbeing. As MZ twins share nearly 100% of their DNA sequence as well as much of their environment during childhood and adolescence, they form the ideal case-control group^34^ as DNA methylation differences that will be found in a discordant EWAS model can be ascribed with more certainty to environmental influences and rule out most SNP-methylation effects. Epigenome-wide association studies using the discordant twin pair design have already been successfully applied for a number of behavioral traits, including schizophrenia and bipolar disorder, aggressive behavior, and autism^37–39^. However, we did not identify significant methylation differences within pairs of discordant twins in a replication analysis of the 10 top CpGs from the meta-analysis, with multiple testing correction for 10 tests. Since MZ pairs who are very discordant for wellbeing are rare, the sample size of this analysis was small and power was limited^40,41^. According to a previously published power analysis^40^, the power to detect DNA methylation level differences with 15 discordant pairs at epigenome-wide significance after multiple testing correction essentially approaches 0. This sample size only has fairly decent power (>=70%) to detect within-pair DNA methylation differences of 10% or larger at nominal significance (alpha 0.05). Such large within-pair differences were not observed among top-ranking sites in the current analysis.

## Strengths and weaknesses

This is the largest EWAS of wellbeing to date. The large sample size was achieved by including multiple measures of satisfaction with life and other wellbeing questionnaires including participants from multiple countries and all ages in a meta-analysis and analyzing DNA methylation data from blood. While this approach maximizes the power to detect common DNA methylation signatures of satisfaction with life and wellbeing, a limitation of this approach is that it will dilute age-and sex specific effects. Furthermore, the power to find statistically significant associations is reduced due to heterogeneity in phenotypic measurements and/or environmental background characteristics, as well as non-identical processing methods of methylation data. Our results indicate that larger EWA studies, or studies focusing on more homogenous measures of wellbeing, are needed to identify DNA methylation sites associated with wellbeing. In addition, other methods such as summary-data-based Mendelian Randomization^42^ can be applied to summary statistics from large GWAS meta-analyses of wellbeing and summary statistics of methylation QTL analyses to increase our understanding of the functional consequences of SNP effects on wellbeing via DNA methylation, and to examine causal relationships between DNA methylation and wellbeing.

## Conclusions

In summary, in this first EWAS meta-analysis of wellbeing to date, we did not detect epigenome-wide significant associations between wellbeing and DNA methylation measured in blood while correcting for typical covariates including white blood cell counts, smoking and BMI. In addition, we found no significant DNA methylation differences in MZ twin pairs highly discordant for wellbeing. Larger meta-analyses and studies with more homogenous wellbeing measures are warranted.

### Ethical Statement

The research was conducted in accordance with the Declaration of Helsinki. Informed consent was obtained from all participants. Cohort-specific details about study approval and consent is provided below.

**ALSPAC:** Ethical approval for the study was obtained from the ALSPAC Ethics and Law Committee and the Local Research Ethics Committees. Informed consent for the use of data collected via questionnaires and clinics was obtained from participants following the recommendations of the ALSPAC Ethics and Law Committee at the time. Consent for biological samples has been collected in accordance with the Human Tissue Act (2004).

**DTR**: For the surveys of the Danish Twin Registry informed consent was obtained from all participants, and the surveys were approved by the Regional Scientific Ethical Committees for Southern Denmark (approvals S-VF-19980072, S-VF-20040241 and S-20090033). This study was approved by the University of Southern Denmark’s Research & Innovation Organization (SDU RIO) under approval 17/57666.

**FTC:** Informed consent was obtained upon contact with the study subjects before questionnaire information was collected, and when clinical investigations were undertaken with sampling of biological material. The ethics approvals for the study were obtained from the ethics committees of the Hospital District of Helsinki and Uusimaa.

**GenR**: The study protocol was approved by the Medical Ethical Committee of the Erasmus Medical Centre, Rotterdam. Written informed consent was obtained for all participants.

**KORA:** The KORA cohort ethical approval was granted by the ethics committee of the Bavarian Medical Association. KORA was carried out in accordance with the principles of the Declaration of Helsinki. This covers consent for the use of biological material, including genetics. All research participants have signed informed consent prior to taking part in any research activities.

**LBC1921:** Ethics permission was obtained from the Lothian Research Ethics Committee and the Scotland A Research Ethics Committee. Participants provided written informed consent. **LBC1936:** Ethics permission was obtained from the Lothian Research Ethics Committee and the Scotland A Research Ethics Committee. Participants provided written informed consent. **LLD:** The LifeLines DEEP study was approved by the ethics committee of the University Medical CenterGroningen, document number METC UMCG LLDEEP: M12.113965. All participants signed an informed consent form before study enrollment. All procedures performed in studies involving human participants were in accordance with the ethical standards of the institutional and/or national research committee and with the 1964 Helsinki declaration and its later amendments or comparable ethical standards.

**NFBC66:** The written informed consents were given by the participants and studies were approved by the Northern Ostrobothnia Hospital District Ethical Committee 94/2011 (12.12.2011).

**NFBC86:** The written informed consents were given by parents and adolescents and studies were approved by the Northern Ostrobothnia Hospital District Ethical Committee 108/2017 (15.1.2018).

**NTR:** Informed consent was obtained from all participants. The study was approved by the Central Ethics Committee on Research Involving Human Subjects of the VU University Medical Centre, Amsterdam, an Institutional Review Board certified by the U.S. Office of Human Research Protections (IRB number IRB00002991 under Federal-wide Assurance-FWA00017598; IRB/institute codes, NTR 03-180).

**NAS**: The Normative Aging Study is overseen by the Institutional Review Board at Veterans Affairs Boston Healthcare System. At each study visit, participants provided written informed consent.

**QIMR:** Informed consent was obtained from all participants. The study was approved by the QIMR Berghofer Human Research Ethics Committee.

## Acknowledgements

**ALSPAC:** ALSPAC is a longitudinal study of pregnant women and their children resident in Avon, UK with expected dates of delivery between 1st April 1991 and 31st December 1992. Of all those invited to take part in the study, 20,248 pregnancies were identified as being eligible and the initial number of pregnancies enrolled was 14,541 for 14,203 unique mothers. When the oldest children were approximately 7 years of age, an attempt was made to bolster the initial sample with eligible cases who had failed to join the study originally. The total sample size for analyses using any data collected after the age of seven is therefore 15,447 pregnancies for a total of 14,833 unique mothers. 12,113 of their partners have been in contact with the study by providing data and/or formally enrolling when this started in 2010. 3,807 partners are currently enrolled.We are extremely grateful to all the families who took part in this study, the midwives for their help in recruiting them, and the whole ALSPAC team, which includes interviewers, computer and laboratory technicians, clerical workers, research scientists, volunteers, managers, receptionists and nurses. Ethical approval for the study was obtained from the ALSPAC Ethics and Law Committee and the Local Research Ethics Committees. Informed consent for the use of data collected via questionnaires and clinics was obtained from participants following the recommendations of the ALSPAC Ethics and Law Committee at the time. Consent for biological samples has been collected in accordance with the Human Tissue Act (2004). Please note that the study website contains details of all the data that is available through a fully searchable data dictionary and variable search tool: http://www.bristol.ac.uk/alspac/researchers/our-data/

**FTC**: We gratefully thank all the participating twins and their families for their long term commitment to the FTC study. We acknowledge the contribution of the study nurses and other staff of the FTC responsible for data collection and management.

**GenR**: The Generation R Study is conducted by Erasmus MC, University Medical Center Rotterdam in close collaboration with the School of Law and Faculty of Social Sciences of the Erasmus University Rotterdam, the Municipal Health Service Rotterdam area, Rotterdam, the Rotterdam Homecare Foundation, Rotterdam and the Stichting Trombosedienst & Artsenlaboratorium Rijnmond (STAR-MDC), Rotterdam. We gratefully acknowledge the contribution of children and parents, general practitioners, hospitals, midwives and pharmacies in Rotterdam. The study protocol was approved by the Medical Ethical Committee of the Erasmus Medical Centre, Rotterdam. Written informed consent was obtained for all participants. The generation and management of the Illumina 450K methylation array data (EWAS data) for the Generation R Study was executed by the Human Genotyping Facility of the Genetic Laboratory of the Department of Internal Medicine, Erasmus MC, the Netherlands. We thank Mr. Michael Verbiest, Ms. Mila Jhamai, Ms. Sarah Higgins, Mr. Marijn Verkerk and Dr. Lisette Stolk for their help in creating the EWAS database. We thank Dr. A. Teumer for his work on the quality control and normalization scripts.

**KORA**: We thank all participants for their long-term commitment to the KORA study, the staff for data collection and research data management and the members of the KORA Study Group (https://www.helmholtz-munich.de/en/epi/cohort/kora) who are responsible for the design and conduct of the study.

**Lothian Birth Cohorts**: We thank the cohort participants and team members who contributed to these studies and Prof Riccardo Marioni for methylation QC.

**NAS:** We would like to thank the participants for their engagement in the NAS over the past 60+ years. We would also like to thank Dr. Amar Mehta, formerly of Harvard School of Public Health, for his assistance in conducting the analyses.

**NFBC:** We thank all cohort members and researchers who have participated in the study. We also acknowledge the work of the NFBC project center.

**NTR**: NTR warmly thanks all participants. Epigenetic data were generated at the Human Genomics Facility (HuGe-F) at ErasmusMC Rotterdam.

**QIMR**: We are grateful for the twins who participated in the study and all those who helped in carrying it out. We thank Ann Eldridge, Marlene Grace, and Kerrie McAloney (sample collection); David Smyth, Harry Beeby, and Daniel Park (IT support); and Anjali Henders and Leanne McNeil (laboratory support).

## Data and code availability

Genome-wide summary statistics from the EWAS meta-analysis are provided in Supplementary Tables 6 (Model 1) and 7 (Model 2). The SOP is available in supplementary information 1. EWAS linear model and gee scripts are provided in supplementary information 2. The meta-analysis script is provided in supplementary information 3.

## Supplementary Files

Supplementary Information 1 – EWAS SOP

Supplementary Information 2 – EWAS linear model and gee scripts (R)

Supplementary Information 3 – Meta-analysis code (METAL)

Supplementary Table 1 – Cohort details

Supplementary Table 2 – Wellbeing descriptives individual cohorts

Supplementary Table 3 – Information methylation data individual cohorts

Supplementary Table 4 – R-code and information about the EWAS analyses

Supplementary Table 5 – Genotype QC information

Supplementary Table 6 - EWAS meta-analysis summary statistics Model 1

Supplementary Table 7 - EWAS meta-analysis summary statistics Model 2

Supplementary Table 8 - Methylation score analysis results

Supplementary Table 9 - MS and PGS results

Supplementary Table 10 - Discordant and concordant monozygotic twin pair analysis

Supplementary Figure 1: Manhattan plot for the EWAS meta-analysis of model 1

Supplementary Figure 2: QQ-Plot for the EWAS meta-analysis of model 1

Supplementary Figure 3: QQ-Plot from the EWAS in wellbeing discordant twin pairs

