## Supplementary Information 3 for "Epigenome-wide association study meta-analysis of wellbeing"

**Supplementary information 3 - METAL meta-analysis code model 1.**

MARKER	cgid

WEIGHT	N

#ALLELE	EFFECT_ALLELE NON_EFFECT_ALLELE

EFFECT	Beta_SWB

STDERR	SE_SWB

PVAL	Pval_SWB

PROCESS Wellbeing_ALSPAC_happygenerally_Model1Ffilteredadjusted.txt

PROCESS Wellbeing_DTR_Happy2_Model1Ffilteredadjusted.txt

PROCESS Wellbeing_FTC2014_Model1Ffilteredadjusted.txt

PROCESS Wellbeing_KORAF4_Model1Ffilteredadjusted.txt

PROCESS Wellbeing_LBC1921_Model1Ffilteredadjusted.txt

PROCESS Wellbeing_LBC1936_Model1Ffilteredadjusted.txt

PROCESS Wellbeing_LLD_Model1Ffilteredadjusted.txt

PROCESS Wellbeing_NFBC1966_Model1Ffilteredadjusted.txt

PROCESS Wellbeing_NFBC1986_Model1Ffilteredadjusted.txt

PROCESS Wellbeing_QIMRB_Model1Ffilteredadjusted.txt

PROCESS Wellbeing_VatNormAge_lsi_Model1Ffilteredadjusted.txt

PROCESS Wellbeing_NTR_Model1Filteredadjusted_JVD.txt

\# Execute meta-analysis

ANALYZE HETEROGENEITY
