## Supplementary Information 2 for "Epigenome-wide association study meta-analysis of wellbeing"

**Supplementary information 2 - EWAS linear model and gee code**

\# 1 linear model code (for unrelated individuals)

#==============================================================================================================================================================================================#

\# CHECKLIST BEFORE DOING YOUR EWAS ANALYSIS

\# 1. Read all comments in this script

\# 2. You have 2 data objects: 1=methylation data. 2=Phenotype/covariate data.

\# 3. Your methylation data is on the beta-value scale (0-1). Bad probes and bad samples have been removed. Data have been normalized (examples of normalization procedures are: Functional Normalization, Quantile Normalization etc).

\# 4. Your methylation data contains numeric values. Missing values are denoted by NA.

\# 5. Your methylation data has the following format: rows=samples, columns= methylation sites.

\# 6. Your methylation data has the following column names: Illumina probe ID (e.g. "cg1234567")

\# 7. Your phenotype/covariate data contains: data on the phenotype and all covariates.

\# 8. Your phenotype and your covariates are numeric values. Missing values in your phenotype and covariates are denoted by NA.

\# 9. Your phenotype/covariate data has the following format: rows=individuals, columns=variables (covariates/phenotype).

\# 10. Your phenotype/covariate data and methylation data have the same number of rows (samples), and are in the exact same order.

\# 11. Your data does not contain: Family members or multiple samples from a single individual. If this situation does occur in your data, please use a different model that allows to correct for clustering of observations (see other example script).

\# NOTE: In the example code, the methylation data object (1) is called "BetaFunNorm", the phenotype/covariate data object (2) is called "mydata". The phenotype (Wellbeing) is called "swb".

#==============================================================================================================================================================================================#

#==============================================================================================================================================================================================#

\# EWAS meta-analysis Aggression/wellbeing	- Example Code Linear Model

\# Example Code for testing the association between Aggression / wellbeing in cohorts that do not include family members and/or twins

\# Author: Jenny van Dongen

\# Date: January-2016

\# DNA methylation level is the outcome, the phenotype (Aggression/wellbeing) is the predictor

\# This example uses a linear regression model, as implemented in the R-package lm

#==============================================================================================================================================================================================#

#==============================================================================================================================================================================================#

\# PLEASE ADAPT THIS PART TO SPECIFY THE VARIABLE LABELS OF THE PHENOTYPE AND COVARIATES IN YOUR DATASET

\# In the code below, you should adapt: covariates, if you would like to include additional/other covariates, or want to adapt to the variable name of your covariates (example: if in your data, the variable age has variable label "AGE", you should change "age" to "AGE" in the code below.

\# In the code below, you should NOT adapt: 1. The first line. 2. the outcome variable (this should be CpGi). 3. data=data

#Model 1

LModel1 <- function (data)

{

lm(CpGi~swb+sex+age+age2+Mono_Perc+Eos_Perc+Neut_Perc +Array_rownum+PC3+PC4+PC5,data=data)

}

#Model 2

LModel1 <- function (data)

{

lm(CpGi~swb+sex+age+age2+Mono_Perc+Eos_Perc+Neut_Perc +Array_rownum+PC3+PC4+PC5+BMI+smoking,data=data)

}

#==============================================================================================================================================================================================#

#==============================================================================================================================================================================================#

\# Script for applying the model specified above to multiple methylation sites, and collecting the output from multiple sites in 1 table.

\# In this example, methylation data are added to the object "mydata" (for 1 CpG at a time), and the linear model is subsequently applied to "mydata".

\# VERY IMPORTANT: Example below assumes that the rows (individuals) in mydata and BetaFunNorm are in the same order.

\# We first apply model 1 to 1 methylation site to obtain the number of estimates (Nestimates) in the output (Nestimates depends on how many covariates you include in the model):

mydata$CpGi <- BetaFunNorm[,1]

coeff <- coef(summary(LModel1(mydata)))

Nestimates <- nrow(coeff)

Nestimates

Ncpgs <-ncol(BetaFunNorm)

Ncpgs

pval <- matrix(NA,Ncpgs,Nestimates)

estimate <- matrix(NA,Ncpgs,Nestimates)

SE <- matrix(NA,Ncpgs,Nestimates)

tval <- matrix(NA,Ncpgs,Nestimates)

N <- matrix(NA,Ncpgs,1)

\# Now we apply model 1 to all methylation sites and extract the output:

for (i in 1:Ncpgs)

{

mydata$CpGi <- BetaFunNorm[,i]

\# N= sample size (probe-specific):

N[i,] <- length(which(!is.na(mydata$CpGi)))

coeff <- coef(summary(LModel1(mydata)))

for (j in 1: Nestimates)

{

estimate[i,j] <- coeff[j,1]

SE[i,j] <- coeff[j,2]

tval[i,j] <- coeff[j,3]

pval[i,j] <- coeff[j,4]

}

}

\# 4 output tables are created: estimate contains the beta-values from the regression model. SE contains the SEs of the beta-values, tval contains the t-values, pval contains the p-values.

\# Format: rows= methylation sites. columns= intercept + all predictors.

\# Below, rownames and columnames are assigned.

\# example: colnames(estimate) <- rownames(coeff). Here: the new column names that are assinged to the output table estimate are: intercept + the names of all predictors in the model (Wellbeing + covariates).

\# rownames(estimate) <- colnames(BetaFunNorm). Here: the new rownames that are assinged to the output table estimate are the illumina probeids (e.g. "cg1234567"). This only works if your methylation data object contains these IDs as column names.

colnames(estimate) <- rownames(coeff)

colnames(SE) <- rownames(coeff)

colnames(tval) <- rownames(coeff)

colnames(pval) <- rownames(coeff)

rownames(estimate) <- colnames(BetaFunNorm)

rownames(SE) <- colnames(BetaFunNorm)

rownames(tval) <- colnames(BetaFunNorm)

rownames(pval) <- colnames(BetaFunNorm)

rownames(N) <- colnames(BetaFunNorm)

\# Please provide the output statistics for the variable Wellbeing and smoking (not for the other covariates).

\# 2 gee model code (for related individuals)

#==============================================================================================================================================================================================#

\# CHECKLIST BEFORE DOING YOUR EWAS ANALYSIS

\# 1. Read all comments in this script

\# 2. You have 2 data objects: 1=methylation data. 2=Phenotype/covariate data.

\# 3. Your methylation data is on the beta-value scale (0-1). Bad probes and bad samples have been removed. Data have been normalized (examples of normalization procedures are: Functional Normalization, Quantile Normalization etc).

\# 4. Your methylation data contains numeric values. Missing values are denoted by NA.

\# 5. Your methylation data has the following format: rows=samples, columns= methylation sites.

\# 6. Your methylation data has the following column names: Illumina probe ID (e.g. "cg1234567")

\# 7. Your phenotype/covariate data contains: data on the phenotype, all covariates, and a family-ID/Twin Pair-ID.

\# 8. Your phenotype and your covariates are numeric values. Missing values in your phenotype and covariates are denoted by NA.

\# 9. Your phenotype/covariate data has the following format: rows=individuals, columns=variables (covariates/phenotype).

\# 10. Your phenotype/covariate data and methylation data have the same number of rows (samples), and are in the exact same order.

\# 11. Your phenotype/covariate data and methylation data are sorted on family-ID/Twin Pair-ID.

\# NOTE: In the example code, the methylation data object (1) is called "BetaFunNorm", the phenotype/covariate data object (2) is called "mydata", the family-ID is called "familynumber", and the phenotype (Wellbeing) is called "swb"

#==============================================================================================================================================================================================#

#==============================================================================================================================================================================================#

\# EWAS meta-analysis Aggression/wellbeing	- Example code

\# Example Code for testing the association between Aggression / wellbeing in cohorts that include family members and/or twins (monozygotic and/or dizygotic)

\# Author: Jenny van Dongen

\# Date: January 2016

\# DNA methylation level is the outcome, the phenotype (Aggression/wellbeing) is the predictor

\# This example uses generalized estimation equation models (gee), as implemented in the R-package gee

\# Within the function gee(), the option "id=" should be used to specify the name of the variable in your data that indicates the clusters of observations (e.g. family-ID or Twin Pair-ID)

\# In the example below, we specificy "id=familynumber", because in the example dataset ("mydata"), there is a column called "familynumber", that indicates the family relationships: all individuals belonging to the same family have the same "familynumber"

#==============================================================================================================================================================================================#

#==============================================================================================================================================================================================#

\# !!!!!!!!! VERY IMPORTANT WHEN USING GEE !!!!!!!!!

\# 1. Make sure that you use the option: corstr="exchangeable"

\# 2. Within the function gee(), specify: id= >familynumber< , where >familynumber< is the variable name of family-ID/Twin Pair-ID in your dataset

\# 3. You MUST sort your data (phenotype AND methylation) on >familynumber< . [If you do not sort on > familynumber <, your analysis will NOT be corrected for clustering within families]

\# 4. From the output of gee(), extract the "robust results": you should calculate the p-value based on the RobustZscore and report the robustSE that is in the output of gee. [If you take the column SE (instead of RobustSE), your results will NOT be corrected for clustering within families]

#==============================================================================================================================================================================================#

install.packages("gee")

library(gee)

?gee

#==============================================================================================================================================================================================#

\# PLEASE ADAPT THIS PART TO SPECIFY THE VARIABLE LABELS OF THE PHENOTYPE, COVARIATES, AND family-ID/Twin Pair-ID IN YOUR DATASET

\# In the code below, you should adapt: 1. id=>familynumber<. 2. covariates, if you would like to include additional/other covariates, or want to adapt to the variable name of your covariates (example: if in your data, the variable age has variable label "AGE", you should change "age" to "AGE" in the code below.

\# In the code below, you should NOT adapt: 1. The first line. 2. the outcome variable (this should be CpGi). 3. family=gaussian (this is the required specification for a contiuous outcome variable). 3. corst=exchangeable. 4. maxiter=100 (do not use a amaller number)

\# model 1

geemodel1 <- function (data)

{

gee(CpGi~swb+sex+age+age2+Mono_Perc+Eos_Perc+Neut_Perc+ Array_rownum+PC3+PC4+PC5,data=data, id=familynumber, family=gaussian, corstr="exchangeable", maxiter=100, na.action=na.omit)

}

\# model 2

geemodel2 <- function (data)

{

gee(CpGi~swb+sex+age+age2+Mono_Perc+Eos_Perc+ Neut_Perc+Array_rownum+PC3+PC4+PC5+BMI+smoking,data=data, id=familynumber, family=gaussian, corstr="exchangeable", maxiter=100, na.action=na.omit)

}

#==============================================================================================================================================================================================#

\# SORTING BY FAMILY-ID

\# Example: To sort the phenotype data on family-ID/Twin Pair-ID (in this example, the family identifier is named "familynumber"):

mydata <- mydata[order(phenodata$familynumber),]

\# note: you should sort your methylation data in the same order

#==============================================================================================================================================================================================#

\# Script for applying the model specified above to multiple methylation sites, and collecting the output from multiple sites in 1 table.

\# In this example, methylation data are added to the object "mydata" (for 1 CpG at a time), and the geemodel is subsequently applied to "mydata".

\# Example below assumes that the rows (individuals) in mydata and BetaFunNorm are in the same order and are sorted on family-ID/Twin Pair-ID.

\# We first apply model 1 to 1 methylation site to obtain the number of estimates (Nestimates) in the output (Nestimates depends on how many covariates you include in the model):

mydata$CpGi <- BetaFunNorm[,1]

coeff <- summary(geemodel1(mydata))$coefficients

Nestimates <- nrow(coeff)

Nestimates

Ncpgs <-ncol(BetaFunNorm)

Ncpgs

pval <- matrix(NA,Ncpgs,Nestimates)

estimate <- matrix(NA,Ncpgs,Nestimates)

RobustSE <- matrix(NA,Ncpgs,Nestimates)

RobustZ <- matrix(NA,Ncpgs,Nestimates)

N <- matrix(NA,Ncpgs,1)

\# Now we apply model 1 to all methylation sites and extract the output:

for (i in 1:Ncpgs)

{

mydata$CpGi <- BetaFunNorm[,i]

\# N= sample size (probe-specific):

N[i,] <- length(which(!is.na(mydata$CpGi)))

coeff <- summary(geemodel1(mydata))$coefficients

for (j in 1: Nestimates)

{

estimate[i,j] <- coeff[j,1]

RobustSE[i,j] <- coeff[j,4]

RobustZ[i,j] <- coeff[j,5]

pval[i,j] <- 2*pnorm(-abs(coeff[j,5]))

}

}

\# It is advisable to check if the above column numbers in the gee ouput (coeff) correspond to the correct columns (in the version of the gee package that you are using).

\# The corrected p-value is computed as follows 2*pnorm(-abs(RobustZvalue))

\# You should do this because gee() itself does not provide the corrected p-value (it provides only the 'Robust' SE and 'Robust' Z-value).

\# Please do NOT extract the Pvalue-column from gee (it is not corrected for family structure).

colnames(estimate) <- rownames(coeff)

colnames(RobustSE) <- rownames(coeff)

colnames(RobustZ) <- rownames(coeff)

colnames(pval) <- rownames(coeff)

rownames(estimate) <- colnames(BetaFunNorm)

rownames(RobustSE) <- colnames(BetaFunNorm)

rownames(RobustZ) <- colnames(BetaFunNorm)

rownames(pval) <- colnames(BetaFunNorm)

rownames(N) <- colnames(BetaFunNorm)

\# 4 output tables are created: estimate contains the beta-values from the regression model. RobustSE contains the Robust SEs of the beta-value, RobustZ contains the Robust Z-values, pval contains the p-values.

\# Format: rows= methylation sites. columns= intercept + all predictors in geemodel (Wellbeing + covariates).

\# Above, rownames and columnames are assigned.

\# example: colnames(estimate) <- rownames(coeff) # Here: the new columnames that are assinged to the output table estimate are: intercept + the names of all predictors in geeemodel.

\# rownames(estimate) <- colnames(BetaFunNorm) # Here: the new rownames that are assinged to the output table estimate are the illumina probeids (e.g. "cg1234567"). This only works if your methylation data object contains these IDs as column names.

\# Please provide the output statistics for the variable Wellbeing and smoking (not for the other covariates).
