## Supplementary Figure 1 for "Epigenome-wide association study meta-analysis of wellbeing"

**Supplementary figures**

**Supplementary Figure 1:** Manhattan plot for the EWAS meta-analysis of model 1

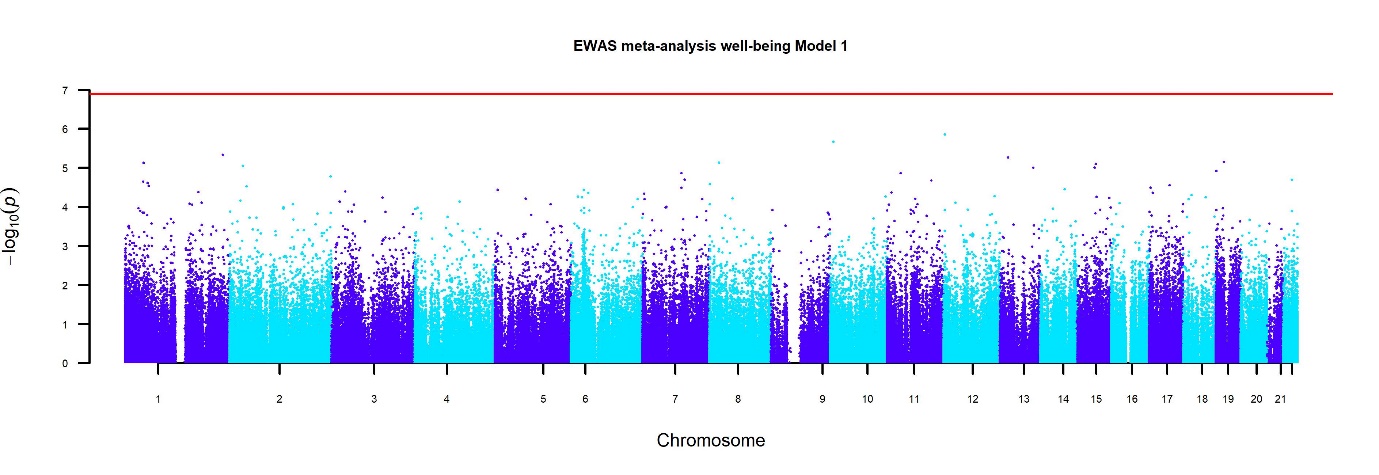

The Manhattan plot shows the p-values for autosomal methylation sites from the EWAS meta-analysis of model 1, without correction for smoking and BMI.

**Supplementary Figure 2:** QQ-Plot for the EWAS meta-analysis of model 1

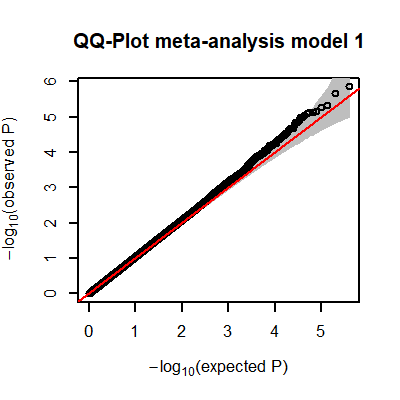

QQ-plot from the EWAS meta-analysis of model 1, without correction for smoking and BMI. The inflation factor (lambda) was 1.03.

**Supplementary Figure 3:** QQ-Plot from the EWAS in wellbeing discordant twin pairs

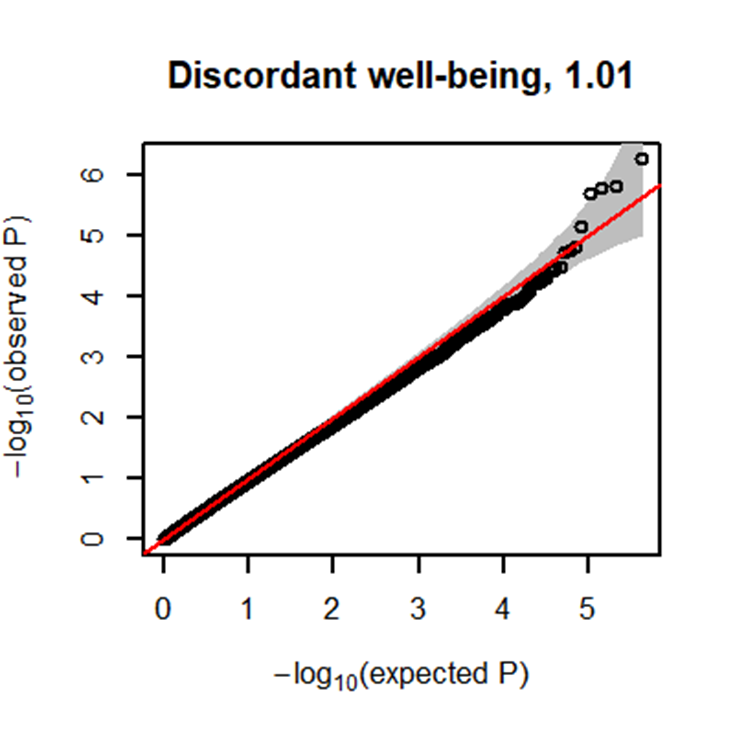

QQ-Plot from the EWAS in wellbeing discordant monozygotic twin pairs. The inflation factor (lambda) was 1.01.
