## Supplementary Information 1 for "Epigenome-wide association study meta-analysis of wellbeing"

**EWAS Wellbeing Analysis Plan**

This document details a standard operating procedure (SOP) for the data analyst that will performing the EWA analyses on wellbeing (WB) in each of the participating cohorts. Standardization of the procedures will increase the precision of the final meta-analyses across all samples of the consortium.

**Contact details**

Questions about this SOP can be directed to:

Bart Baselmans,

Jenny van Dongen,

Meike Bartels,

*Please, for clarity, CC both contact persons in your email.*

**Background**

The goal of this initiative is to carry out a large-scale Epigentic-Wide Association Analysis (EWAS) meta-analysis on wellbeing (WB). The phenotype is defined broadly to include all positively framed subjective assessments of WB. WB is an increasingly major research topic in both (biological) psychology (e.g. Bartels et al., 2015) and in economics (e.g. Graham 2010). There is much evidence that WB affect immune function and health status (Marsland, Pressman & Cohen, 2007), as well as other life outcomes. There is much interest in the genetic basis of WB, with twin studies suggesting it is ~40% heritable (Bartels, 2015). Additionally, a recently performed GWAS meta-analysis involving 300K participants, found the first genetic variants significantly associated with WB (Okbay et al., 2016). However, from these studies we know that besides genotype, environmental factors play an important role in explaining the variance in WB. Epigenetic mechanisms, such as DNA methylation, regulate gene expression, and may mediate genetic and environmental effects on WB. A recently performed EWAS (N = ~2500) on WB on whole blood Illumina 450K methylation data, collected at the Netherlands Twin Register (NTR) found the first evidence for significant associations between WB and DNA methylation (Baselmans et al., 2015). As more cohorts have available methylation data, as well as WB data, we now propose to perform an EWAS meta-analyis of WB. We would like to ask each cohort to perform the EWAS analysis on their cohort and to provide the results (summary statistics) to us for the meta-analysis.

**Epigenetic clock**

In addition to the EWAS approach, we aim to test the relationship between wellbeing and the epigenetic clock (DNA methylation age, DNAmAge). Several studies suggest a U-shaped relationship between SWB and age, with the lowest point approximately in midlife (Blanchflower & Oswald, 2008; Layard, 2010). We hypothesize that this U-shaped relationship has a biological mechanisms, which might be reflected in the epigenetic clock.

**Timeline**

We would like to ask you to send us your EWAS results by March 14. Depending on the meta-analysis results, we plan to perform secondary analysis for top its, including analyses to test if DNA methylation level is associated with gene expression level (RNA seq and/or Affymetrix U219 array). If you have gene expression data an would like to contribute o the secondary analysis stage, or if you have andy ideas for interesting follow-up analyses, please let us know.

**Help**

Please let us know if there is anything we can do to help you, or if you have any comments or suggestion, by e-mailing us:

Bart Baselmans,

Jenny van Dongen,

Meike Bartels,

***Multiple measures of Wellbeing or DNA methylation***

*If you have data on WB available from multiple measurements instruments or collected at different ages, or if you have DNA methylation data available from samples collected at multiple ages or from multiple tissues. Please let us know so we can discuss the optimal analysis strategy.*

**Phenotype Definition**

While there are distinctions between various facets of WB (e.g., Seligman, 2011), measures of different facets are correlated with each other. Moreover, different measures load on a common genetic factor (Bartels & Boomsma, 2009). For this reason, and to maximize sample size, our primary phenotype definition will encompass a broad array of WB measures, treating them as different measures of the same underlying construct. Across different facets of WB, the most common distinction is between “positive affect and “life satisfaction” (e.g., Kahneman & Riis, 2005; Kahneman & Deaton, 2010). Positive affect refers to the frequency and intensity of positive emotions and feeling happy. Typical survey questions that gauge positive affect include “During the past week, I was happy?” and “How would you rate your emotional wellbeing at present?” Life Satisfaction refers to a longer. term evaluation of one’s life. A typical survey question would be “How satisfied are you with your life as a whole?” Positive affect and life satisfaction correlate strongly but show varying salience in different circumstances of people’s lives. Hence, if we find a Epigenetic-wide significant association with the broad WB variable, which pools across different measures, then we will also conduct separate meta-analyses on these two facets of WB in order to understand whether or not this association is specific to positive affect or life satisfaction.

**In sum, the meta-analysis of Well Being will primarily analyze a variety of measures of WB pooled. In follow-up analyses, we may separately study positive affect measures only and life satisfaction measures only.**

**Note:** Cohorts do not need to categorize their own WB measure as a positive affect

measure, a	life	satisfaction	measure, or	neither. Based on the cohort’s uploaded descriptives, which includes the survey question texts, we will make this determination and include some measures but not others in the follow-up analyses.

**Analysis Plan – Summary**

| Methylation data | Illumina 450k array, beta-values, after QC and normalization |
| --- | --- |
| Tissue | We assume that you extracted DNA from peripheral whole blood |
| Phenotype | Wellbeing, on a continuous scale (**higher score should**  **correspond to a higher level of wellbeing**) |
| EWAS analysis | For each methylation site, we test if DNA methylation level is associated with wellbeing score. Methylation level is the outcome, wellbeing is predictor. This test is repeated for all genome-wide methylation sites (that survived cohort-specific methylation data QC). |
| Covariates - EWAS model 1 | Age, Age^2^ sex, white blood cell counts, technical and cohort-specific covariates |
| Covariates - EWAS model 2 | Covariates model 1 + BMI and smoking status |
| EWAS results files | Wellbeing_Cohort_Model1_YYYYMMDD_InitialsAnalyst.csv [5 columns]  Wellbeing_Cohort_Model2_YYYYMMDD_InitialsAnalyst.csv [8 columns]  Requested information: cgid, N, Beta, SE, Pval |
| DNAmAge results file | Wellbeing_CohortName_DNAmAge _YYYYMMDD_InitialsAnalyst.csv  Requested information: Measure, N, Beta, SE, Pval [5 columns] |
| Cohort information file | Wellbeing_Cohort_CohortInformation_YYYYMMDD_InitialsAnalyst.xls |

**Analysis Plan – Details (Three steps)**

***1. Before the analysis: input data for the EWAS and Quality Control***

**DNA methylation**

We will use genome-wide DNA methylation data (illumina 450k array) for the current EWAS. Before running the EWAS, please apply quality control (QC) and normalization. Let us know if we can help you with this. Please perform the analysis on the methylation beta-values (which range from 0 to 1).

**DNA methylation quality control**

Please let us know which QC-filters you have applied to probes and samples and which normalization method you applied to the methylation data you used as input for this EWAS analysis, by completing the excel sheet (Wellbeing_CohortName_CohortInformation_YYYYMMDD_InitialsAnalyst.xls).

We assume that you have removed bad quality samples, and have set probes to NA within a sample (prior to the analysis) based on:

* detection p-value (we advise to set probes with p > 0.01 to NA)

* low bead count on array (we advise to set probes with bead count < 3 to NA)

* raw intensity value of exactly zero (probe not present on array)

* and removed probes with low success rate (we advise to remove probes if success rate < 0.95)

Apart from probes that fail QC in your cohort, please analyze **ALL** genome-wide CpGs (if possible).

Note: We will filter out cross-reactive probes, and probes that harbor a SNP prior to the meta-analysis.

**Phenotype: Wellbeing**

We aim to perform EWAS analyses on a quantitative measure of wellbeing (**Higher score should correspond to a higher level of wellbeing)**. Please let us know what measure you used, by completing the excel sheet (Wellbeing_CohortName_Cohortinformation_YYYYMMDD_InitialsAnalyst.xls).

***2. Analysis***

***2.1: EWAS:***

**Model:**

For all methylation sites, please run 2 models:

1. Methylation ~ Wellbeing + Age +Age^2^ + Sex + WBC percentages + Technical + Cohort Specific

2. Methylation ~ Wellbeing + Age + Sex+ Age^2^ + WBC percentages + Technical + Cohort Specific + BMI +

Smoking

*Note: If your cohort includes related individuals (e.g. family members, twins), please apply a statistical approach that takes the clustering of data into account (e.g. gee or linear mixed models).*

**Covariates**

Age= Age when DNA sample was collected

Age^2^ = standardize age variable before square it

WBC= White blood cell percentage in the same blood sample from which DNA was extracted. If you did not measure white blood percentages in the same sample as used for the DNA methylation measurement, please estimate WBC percentages using a prediction method (e.g. Houseman’s reference based method). For computational reasons, please do not include multiple WBC that are highly correlated with each other or that show very little variation between people in your cohort. For example, in the NTR, we use the following WBC as covariates: monocyte percentage, eosinophil percentage, neutrophil percentage.

Technical covariates + Cohort Specific covariates. Please correct for technical (batch) covariates and other cohort-specific covariates as you deem necessary. For example, at the NTR, we include 450k array row and either sample plate or (carefully chosen) principal components from the methylation data. If your cohort included multiple population ancestries, please take this into account as you think it is most appropriate for your cohort (possible strategies include exclusion of small groups of ‘ethnic outliers’, running the analysis separately by ethnicity or inclusion of principal components based on genotype data).

BMI= Body mass index

Smoking= Smoking status at the moment of blood sampling, 3 levels: 0=never smoked, 1=former smoker, 2=current smoker.

**Software and example R-script**

Feel free to use your own analysis pipeline or preferred software to run the models outlined above. Example R-code with instructions is provided in:

Example LM script_012016.r	[suited for cohorts that include unrelated subjects]

Example gee script_012016.r	[suited for cohorts that include related subjects (family members)]

*2. Analysis*

***2.2 – DNAm age acceleration and wellbeing***

**Estimating DNAm age**

DNAm age acceleration can be easily calculated with the convenient online tool from Steve Horvath: (<http://labs.genetics.ucla.edu/horvath/dnamage/>). Please follow the instructions on the website.

In short:

* Upload raw (un-normalized) methylation beta-values (you may preselect probes or include all genome-wide data)

* Remove the bad quality samples you also removed from your EWAS

* Include at least the probes listed in “datMiniAnnotiation.csv “ (downloadable from the website)

* Upload a sample annotation file, with the column names “SampleID”, “Age”, “Female”, “Tissue”

* Column “Female” should contain 1 for females, and 0 for males.

* Crucial: Methylation file and sample annotation file must be in the same order

* For very large cohorts, it is recommended to upload your data in batches of up to around 1000 samples (and to apply preselection of probes)

* Max file size that can be uploaded is 800 Mb

* Crucial: select the options **Normalize Data** and **Advanced Analysis of Blood**

* Crucial: **do NOT use** the option Fast Imputation

* Hint: First test if your input files are in the correct format by running the calculator without the option normalize data and without the option advanced analysis of blood, which should give you the output very quickly. If you do not receive an email with results within one hour, the format of your input files is not correct.

**Variables**

Please collect the following variables from the DNAm age calculator output file:

* AgeAccelerationResidual

* AHOAdjCellCounts

* AAHAAdjCellCounts

**Model**

Please run 3 models:

1. AgeAccelerationResidual	~ Wellbeing + Sex + BMI + Smoking + Cohort Specific

2. AHOAdjCellCounts	~ Wellbeing + Sex + BMI + Smoking + Cohort Specific

3. AAHAAdjCellCounts	~ Wellbeing + Sex + BMI + Smoking + Cohort Specific

*Note: If your cohort includes related individuals (e.g. family members, twins), please apply a statistical approach that takes the clustering of data into account (e.g. gee or linear mixed models).*

*3. Please send us the following files*

**Cohort information file**

File name:

Wellbeing_CohortName_CohortInformation_YYYYMMDD_InitialsAnalyst.xls

example: Wellbeing_NTR_ CohortInformation _20160119_BML.xls.

***Please complete the 3 sheets of this excel file.***

**EWAS output**

File format: .csv (comma-delimited file)

File name:

Wellbeing_CohortName_Model1_YYYYMMDD_InitialsAnalyst.csv	[5 columns]

Wellbeing_CohortName_Model2_YYYYMMDD_InitialsAnalyst.csv	[8 columns]

example: Wellbeing_NTR_Model1_20160119_BML.csv.

Requested columns (include the headers in your file) **model 1**:

**cgid**	Illumina probe ID

**N**	Sample size

**Beta_SWB**	Effect size (for wellbeing)

**SE_SWB**	Standard error of beta (for wellbeing)

**Pval_SWB**	P-value (for wellbeing)

** We would appreciate it if you could adhere precisely to the column naming scheme as indicated (“Pval_SWB”, not: “pval_swb” or “P”)*

Requested columns (include the headers in your file) **model 2**:

**cgid** Illumina probe ID

**N** Sample size

**Beta_SWB**	Effect size (for wellbeing)

**SE_SWB**	Standard error of beta (for wellbeing)

**Pval_SWB**	P-value (for wellbeing)

**Beta_Smoking**	Effect size (for smoking)

**SE_Smoking**	Standard error of beta (for smoking)

**Pval_Smoking**	P-value (for smoking)

* We will use the summary statistics of the variable smoking for QC purposes.

** We would appreciate it if you could adhere precisely to the column naming scheme as indicated (“Pval_SWB”, not: “pval_swb” or “P”)*

**DNAmAge output**

File format: .csv (comma-delimited file)

File name:

Wellbeing_CohortName_DNAmAge_YYYYMMDD_InitialsAnalyst.csv	[5 columns]

example: Wellbeing_NTR_ DNAmAge_20160119_BML.csv.

Number of rows:4.

Row 1=header

Row 2-4= the 3 DNAmAge acceleration measures

Requested columns (include the headers in your file):

**Measure**	“AgeAccelerationResidual” or **“**AHOAdjCellCounts” or “AAHAAdjCellCounts”

**N**	Sample size

**Beta_SWB**	Effect size (for wellbeing)

**SE_SWB**	Standard error of beta (for wellbeing)

**Pval_SWB**	P-value (for wellbeing)

** We would appreciate it if you could adhere precisely to the column naming scheme as indicated (“Pval_SWB”, not: “pval_swb” or “P”) and use exactly the same names for the DNAmAge acceleration measures to avoid confusion.*

**File upload**

4 files Wellbeing_CohortName_CohortInformation_YYYYMMDD_InitialsAnalyst.xls

Wellbeing_CohortName_Model1_YYYYMMDD_InitialsAnalyst.csv

Wellbeing_CohortName_Model2_YYYYMMDD_InitialsAnalyst.csv

Wellbeing_CohortName_DNAmAge_YYYYMMDD_InitialsAnalyst.

**SFTP server adress:	lisa.surfsara.nl**

**Username:	swb_ewas**

**Password:	SWBmeta2016**

***Please upload your files to the folder /DATA_UPLOAD***

**Thank you for your contribution to this project!**

We realize that filling out excel sheets is not your favorite job. But please note that all the information we are collecting will be used to make sure that the analyses of all cohorts are harmonized before we do the meta-analysis. The success and progress of the meta-analysis crucially depends on the availability of all information, so we would really appreciate it if everyone could return all completed files together with their EWAS results. To put all this information together and run the analysis, we think that 6 weeks should be an appropriate timeframe (assuming that your methylation data is ready to analyze), however, if this is too soon for you, just let us know. Your effort in carefully performing the analysis and completing all information is more important to us than the deadline.

Thank you very much in advance for your cooperation!
